# Beyond stability constraints: a biophysical model of enzyme evolution with selection for stability and activity

**DOI:** 10.1101/399154

**Authors:** Julian Echave

## Abstract

Proteins trace trajectories in sequence space as their amino acids become substituted by other amino acids. The number of substitutions per unit time, the rate of evolution, varies among sites because of biophysical constraints. Several properties that characterize sites’ local environments have been proposed as biophysical determinants of site-specific evolutionary rates. Thus, rate increases with increasing solvent exposure, increasing flexibility, and decreasing local packing density. For enzymes, rate increases also with increasing distance from the protein’s active residues, presumably due to functional constraints. The dependence of rates on solvent accessibility, packing density, and flexibility has been mechanistically explained in terms of selection for stability. However, as I show here, a stability-based model fails to reproduce the observed rate-distance dependence, overestimating rates close to the active residues and underestimating rates of distant sites. Here, I pose a new biophysical model of enzyme evolution with selection for stability and activity (*M*_SA_) and compare it with a stability-based counterpart (*M*_S_). Testing these models on a structurally and functionally diverse dataset of monomeric enzymes, I found that *M*_SA_ fits observed rates better than *M*_S_ for most proteins. While both models reproduce the observed dependence of rates on solvent accessibility, packing, and flexibility, *M*_S__A_ fits these dependencies somewhat better. Importantly, while *M*_S_ fails to reproduce the dependence of rates on distance from the active residues, *M*_SA_ accounts for the rate-distance dependence quantitatively. Thus, the variation of evolutionary rate among enzyme sites is mechanistically underpinned by natural selection for both stability and activity.

## INTRODUCTION

From the early days of Molecular Evolution we know that some protein sites evolve more slowly than others. Ever since, we have interpreted such variation of rates among sites in terms of purifying selection to maintain function and structure: slowly evolving sites are those where mutations are more likely to be discarded by natural selection because they perturb the protein’s structure or function too much [30, 47].

Further insight into the determinants of evolutionary rates came from quantitative studies of the dependence of rates on various metrics that describe the local micro-environment of protein sites. The two metrics that best correlate with rates are *RSA* [10, 16, 17, 48, 50, 61, 62] and *WCN* [16, 35, 52, 54, 61, 62]. *RSA* is the Relative Solvent Accessibility, which measures solvent exposure; *WCN* is the Weighted Contact Number, which measures local packing density. Beyond properties of the static native structure, some work has studied the dependence of rates on dynamical properties such the B-factor (*B*) of X-ray experiments, which measures local flexibility [25, 32, 33, 36, 46, 53]. Evolutionary rate increases with increasing *RSA* and *B* and decreases with increasing *WCN*. These findings have led to the view that the rate of evolution is mainly determined by protein structure, increasing from a slowly evolving, buried, tightly packed, and rigid protein core, towards a rapidly evolving, solvent-exposed, loosely packed, flexible surface [15].

In the previous view, functional constraints play only a minor role, affecting the evolutionary conservation of just a few sites [15], such as the catalytic residues of enzymes and some of their immediate neighbors [3, 59]. However, such a short-range effect of active sites on evolutionary rates seems to be at odds with directed evolution experiments, in which often 2nd or 3rd shell mutations are needed to optimize an enzyme’s function [9, 22]. Also, the very existence of allosteric enzymes, whose activities change due to coupling between distant sites, demonstrates the possibility of long-range couplings [7, 9]. Consistent with this, an early study of *α/β -* barrel enzymes found that evolutionary rate correlates not only with the structural trait *RSA*, but also with distance from the closest active residue *d*_active_ [10]. This finding was confirmed by a recent systematic study of a large and diverse dataset of enzymes that showed that, on average, site-specific substitution rates increase slowly with increasing *d*_active_, which led to the conclusion that active sites influence evolutionary rates at long distances, affecting most protein sites [26].

Studying the dependence of evolutionary rate on metrics such as *RSA, WCN*, *B*, and *d*_active_ is useful to develop phenomenological models which can be used for rate prediction. Moreover, such studies provide some insight into the possible biophysical forces that shape evolutionary sequence divergence. However, as good as they may be for predictive or heuristic purposes, phenomenological models have limited explanatory power. Explaining the origin of rate variation among sites and the relationship between rate and structural and dynamical metrics requires theoretical mechanistic models grounded on first principles.

Several theoretical biophysical models have been used to study evolutionary constraints on sequence divergence [13]. Most of these models assume that protein fitness depends only on protein stability. Such stability-based models account for much of the observed variation of evolutionary rates among sites [14, 24, 25, 27, 35]. Moreover, stability-based models reproduce quite well the dependence of site-specific rates on *RSA, WCN*, and *B* [25, 35]. Thus these traits can largely be considered as proxies of protein stability, which would be the actual underlying cause of differential conservation of protein sites [15].

Even though selection for stability explains the dependence of rates on *RSA, WCN*, and *B*, it is unlikely to account for the observed rate-*d*_active_ dependence, which is probably strongly influenced by selection on activity. The very few studies that consider selection on activity focus on ligand binding [13]. Ligand-binding constraints may lead to longrange epistasis [45], which is consistent with the rate-*d*_active_ dependence found for enzymes. However, rather than ligand binding, the key to enzymatic activity is lowering the activation energy barrier of the catalyzed reaction. For a proper understanding of the dependence of site-specific evolutionary rate on *d*_active_, we need biophysical models that include activity constraints based on realistic principles of enzymatic catalysis.

Here I propose a mechanistic biophysical *stability-activity model* of enzyme evolution, *M*_SA_, that includes selection for stability and activity explicitly. I compare this model with a *stability model, M*_S_, developed previously [14]. Using a diverse dataset of 160 monomeric enzymes, I assess whether *M*_S_ and/or *M*_SA_ account for the observed rate variation among sites and, especially, whether they explain the dependence of rate on the site-specific traits *RSA, WCN*, *B*, and *d*_active_. I will show that *M*_SA_ is the best model, which means that activity constraints influence the pattern of rate variation among sites.

## NEW APPROACH

I propose a new model of enzyme evolution, a stability-activity model, *M*_SA_, and, for the sake of comparison, I also consider a stability-based model *M*_S_. *M*_S_ and *M*_SA_ consider evolution as an origin-fixation process in which at each evolutionary time step a mutant is introduced by random mutation and is fixed, replacing its parent, with a certain fixation probability that accounts for natural selection. In general, any mutation may affect the enzyme’s stability, measured by the change in free energy ΔΔ*G*, and its activity, which I measure by a change in activation free energy ΔΔ*G*^*\**^. In *M*_S_ the fixation probability depends exclusively on ΔΔ*G* and has a single parameter, *a*_S_, that represents selection pressure (Materials and Methods, Eq. 1). For *M*_SA_, the fixation probability depends on ΔΔ*G* and on ΔΔ*G*^*\**^, and on two selection-pressure parameters, *a*_S_ and *a*_A_ (Materials and Methods, Eq. 2).

To calculate ΔΔ*G* and ΔΔ*G*^*\**^, I used the Linearly Forced Elastic Network Model (LFENM), which represents a given protein as an elastic network of nodes (amino acids) connected by harmonic springs (interactions) and models mutations as random perturbations of the lengths of the springs that connect the mutated site to other sites [11, 12, 25, 35]. The mutational stability change, ΔΔ*G*, results from the mechanical stress introduced by the mutation minus the relaxation of part of this stress by modification of the mutant’s conformation (Materials and Methods, Eq. 8). I calculated ΔΔ*G*^*\**^ as the energy necessary to deform the mutant’s catalytic residues so that they adopt their wild type conformation (Materials and Methods, Eq. 9).

Theoretical site-specific rates for model *M*_SA_ for a given protein are calculated as follows. First, an *in silico* mutational scan is performed introducing random mutations at each site and calculating ΔΔ*G* and ΔΔ*G*^*\**^. Then, site-specific rates are calculated using Eq. 12, Eq. 13 and Eq. 2. These depend on the model parameters *a*_S_ and *a*_A_, which represent the degree of selection for stability and activity, respectively, and are chosen to minimize the Mean Square Error between model rates 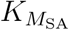 and the empirical rates *K*_obs_. For model *M*_S_ a similar procedure is followed using only ΔΔ*G* and optimizing *a*_S_ to minimize the MSE between 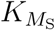 and *K*_obs_. All rates are relative to their protein average.

For a more detailed description of the models see Materials and Methods.

## RESULTS

I applied *M*_S_ and *M*_SA_ to a dataset of 160 monomeric enzymes with diverse sizes, functions, and structures. Since different proteins may be subject to different degrees of selection for stability and/or activity, I performed a protein by protein analysis. For each protein, I performed a mutational scan to calculate ΔΔ*G* and ΔΔ*G*^*\**^ for each of 19 point mutations at each protein site and I calculated the site-specific rates predicted by the models, 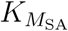 and 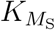, by fitting the models’ parameters to observed rate estimates, *K*_obs_, obtained from sequence alignments using the program `rate4site` [37]. See Supplementary Material file dataset.pdf for details on the proteins studied, their sizes, structural and functional classes, and the parameters of models *M*_SA_ and *M*_S_.

To test the models, I investigated two issues. First, which of the model rates, 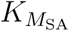 or 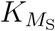, provides a better fit to empirical rates, *K*_obs_. Second, which model provides a better account of the the dependence of observed site-specific rates on the molecular traits *RSA, WCN*, *B*, and *d*_active_.

### Fit between predicted and observed rates: *M*_SA_ outperforms *M*_S_

So far, most biophysical studies have assumed that stability is the main, if not only, determinant of site-specific evolution, disregarding specific activity constraints altogether. Thus the first issue to consider is whether it is at all necessary to add selection for activity explicitly. In other words, are observed rates fit better by *M*_SA_ than by *M*_S_?

Figure 1 shows empirical rates vs. model predictions for protein 1PMI, which I use as an example. (1PMI is the pdb code of the Phosphomannose isomerase of Candida albicans.) Typically, models are compared using *R*^2^, the square Pearson correlation coefficient between predicted and observed rates. For 1PMI, 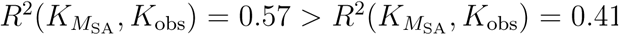. Thus, in this case adding activity constraints increases the explained variance by 16%. I consider *R*^2^ for the sake of comparison with related work and to have an idea of the magnitude of improvement that results from adding activity constraints. However, for nonlinear models, the Akaike Information Criterion (*AIC*) is better for model selection[55]. The best model is that with lowest *AIC*. For 1PMI, Δ*AIC* = *AIC*(*M*_SA_) *-AIC*(*M*_S_) = *-*127.4, thus *M*_SA_ should be selected. Using Δ*AIC*, we can calculate Akaike Weights, *w*(*M*_SA_) and *w*(*M*_S_), which give the weight of evidence for each model. For 1PMI, *w*(*M*_SA_) = 1 and *w*(*M*_S_) = 0. To summarize, for 1PMI, rates predicted with *M*_SA_ fit observed rates better than rates predicted by *M*_S_.

**FIG. 1.**
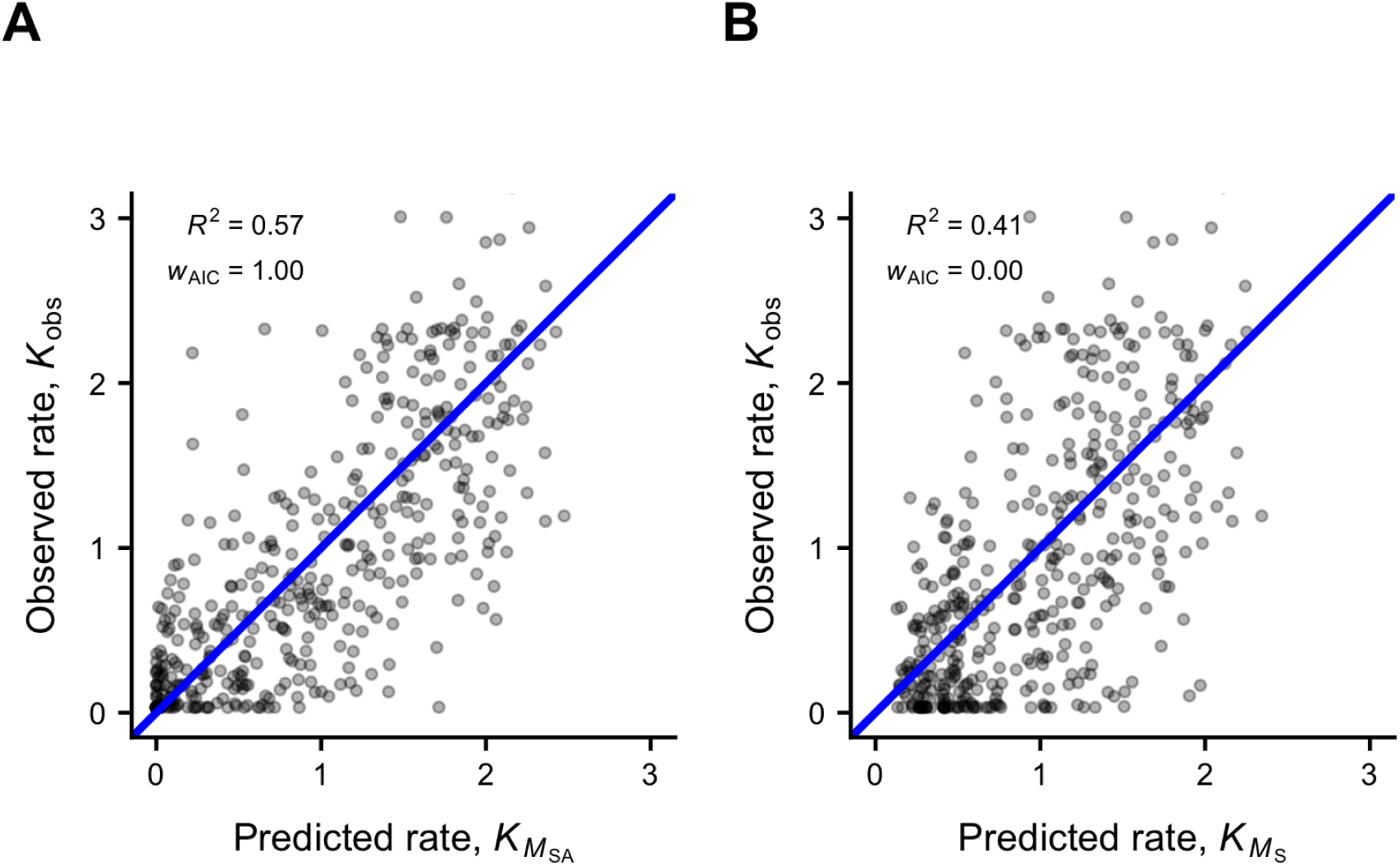
Adding activity constraints improves the fit between observed and predicted rates for 1PMI. Observed site-specific substitution rates for 1PMI vs predictions of the stability-activity model MSA (panel A), and the stability model MS (panel B). Each point represents one protein site. Observed site-specific rates were obtained from multiple sequence alignments using the program `rate4site`. The square correlation coefficients are 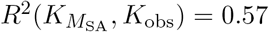 and 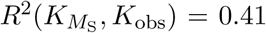. The best model, according to the Akaike Information Criterion, is *M*_SA_, with weight *w*(*M*_SA_) = 1.00.

I repeated the previous analysis for each of the enzymes of the dataset. Figure 2 shows that *M*_SA_ outperforms *M*_S_ for most proteins: *M*_SA_ has the lowest AIC for 131/160 proteins (Binomial test: *p* = 82%, *P* = 6 *×* 10^−17^). Also, *R*^2^(*MSA*) *> R*^2^(*MS*): *<* Δ*R*^2^ *>*=*< R*^2^(*MSA*) *-R*^2^(*MS*) *>*= 0.042 *±* 0.06 (*±* two standard deviations) (t Test: *t* = 12.9, *P* = 0). Thus, adding activity constraints results in better predictions, which means that activity constraints have observable effects on the patterns of rate variation among sites for most proteins.

**FIG. 2.**
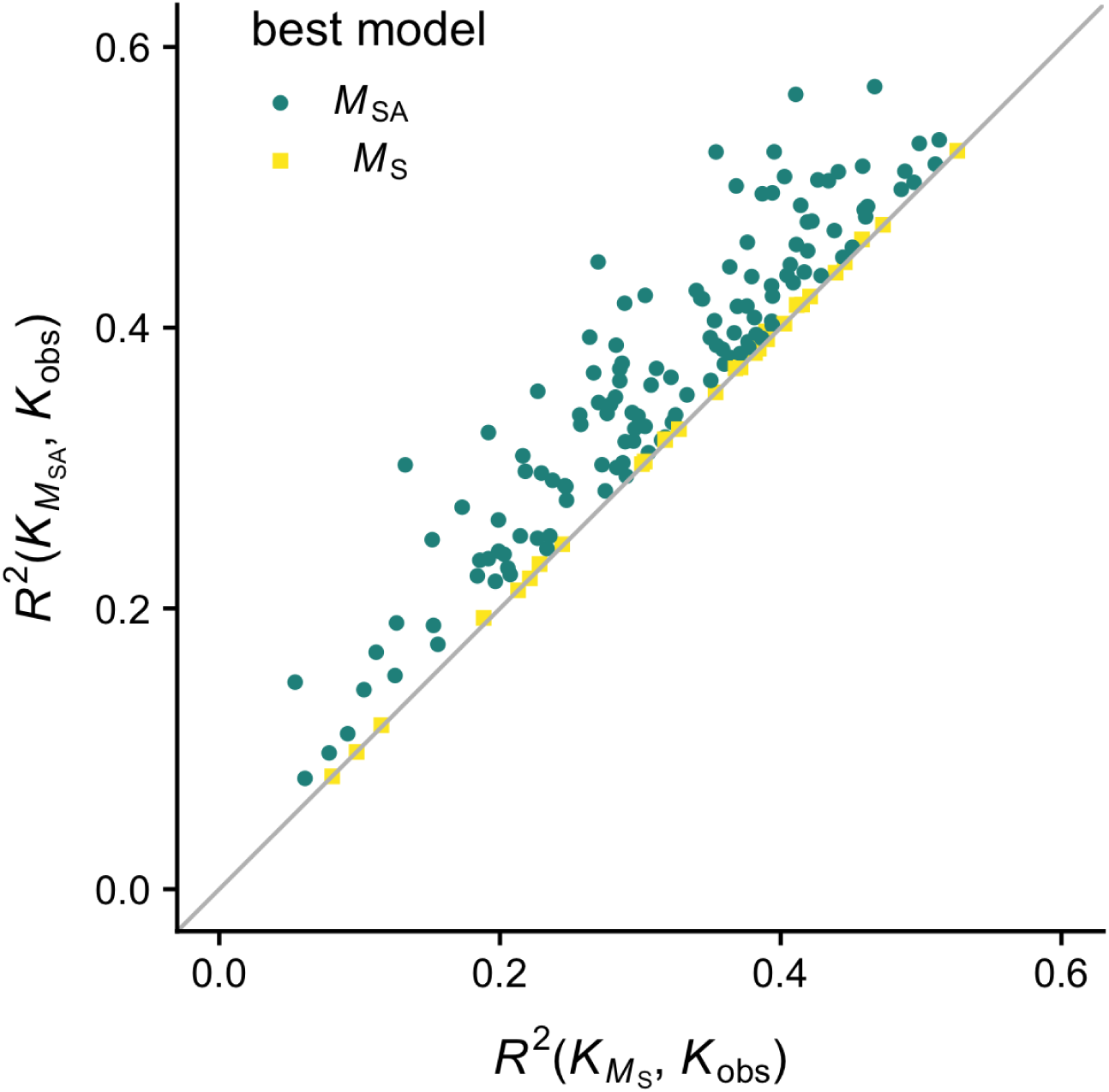
MSA outperforms MS for most proteins. For each protein and each model, goodness of fit is quantified using the square observed-predicted correlation coefficient *R*^2^ and Akaike weights. This figure shows *R*^2^ of model *M*_SA_ vs. *R*^2^ of model *M*_S_. The *y* = *x* line is shown for reference. Colors and shapes indicate the best model according to Akaike weights (i.e. which weights is larger, *w*(*M*_SA_) or *w*(*M*_S_)). For most proteins (131 out of the 160) *M*_SA_ is the best model (*w*(*M*_SA_) *>* 0.5).

### Dependence of site-specific rates on molecular traits: MSA outperforms MS

Beyond the overall goodness of fit, studied in the previous section, a good model should also reproduce how site-specific rates depend on the traits with which they correlate: *RSA, WCN*, *RSA*, and *d*_active_, which I will refer to as molecular traits or empirical predictors. In this section, I explore whether the models account for the dependence of observed rates on these empirical predictors.

Following the scheme of the previous section, I start with the example protein 1PMI. Figure 3 shows observed and model substitution rates vs. the various empirical predictors. For a given *K* (observed or model rate) vs. *X* (empirical predictor), let 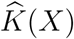 be the smooth fit to the scatter plot obtained using local polynomial regression (shown with lines in Figure 3, see Materials and Methods for details). Let *RMSE*(*M, X*) be the Root Mean Square Error between smCooth trends 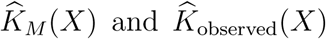, and let Δ*RMSE*(*X*) = *RMSE*(*M*_SA_, *X*) *- RMSE*(*M*_S_, *X*). The model that has lowest *RMSE*(*X*) reproduces better the 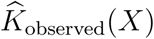 trend. For 1PMI, Δ*RMSE*(*RSA*) = 0.00, Δ*RMSE*(*WCN*) = −0.08, Δ*RMSE*(*B*) = −0.06, and Δ*RMSE*(*d*_active_) = −0.22. Thus, in this case, both models reproduce dependence of rates on *RSA* similarly, *M*_SA_ provides a better fit for the the dependence of rates on *WCN* and *B*, and *M*_SA_ provides a much better account of the dependence of rates on *d*_active_. From Figure 3 we see that relative to *M*_SA_, *M*_S_ overestimates (underestimates) rates at small (large) *d*_active_, small (large) *B* and large (small) *WCN*.

**FIG. 3.**
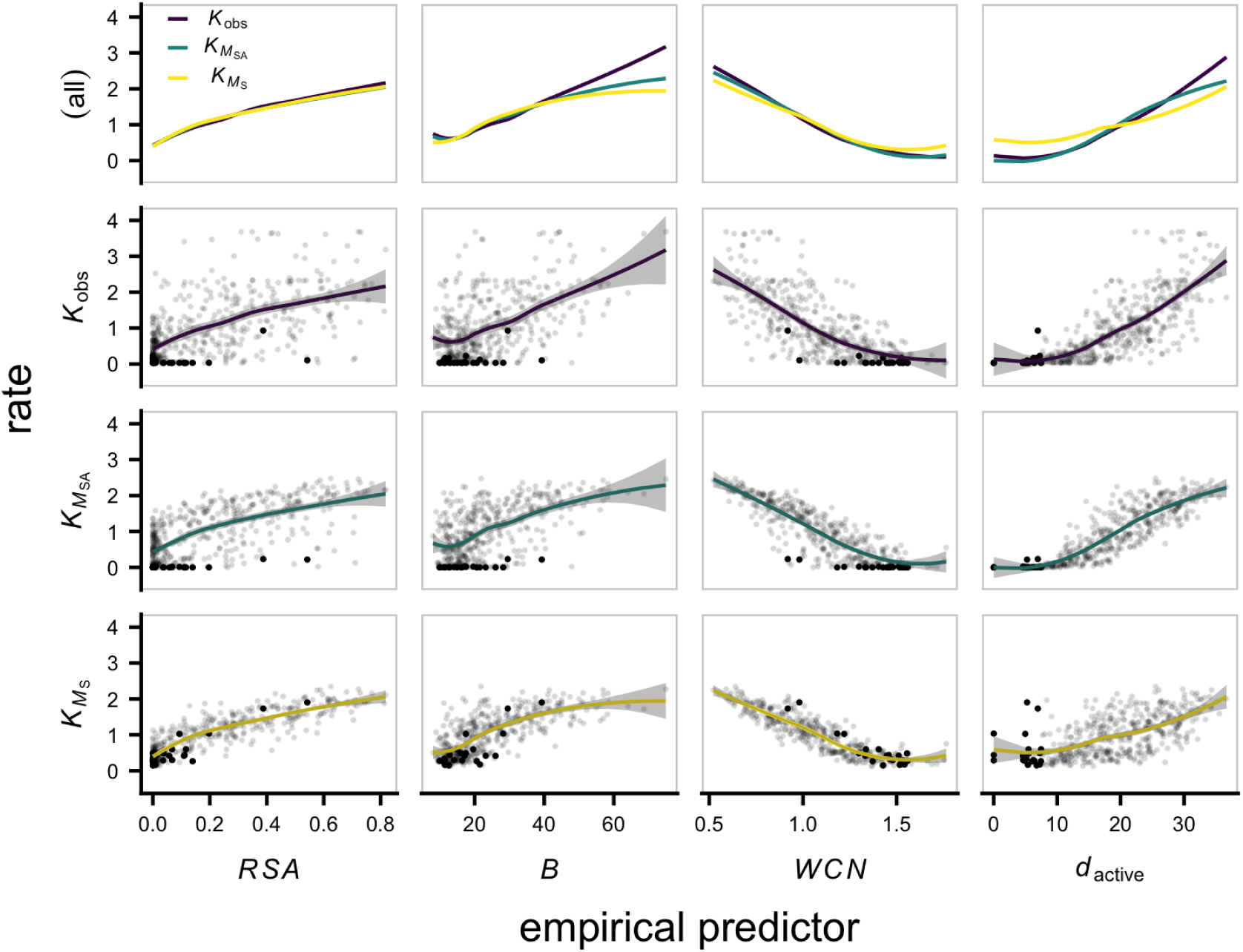
*M*_SA_ provides the best fit of the dependence of site-specific rates on their structural, dynamical, and functional empirical predictors for 1PMI. This figure shows observed and predicted rates vs. four molecular traits that have been found to correlate with rates: *RSA* (relative solvent accessibility), *B* (X-ray B-factor), *WCN* (weighted contact number), and *d*_active_ (distance from closest active residue). The first row shows smooth trends, omitting sites. Rows 2-4 show the smooth fits together with actual rates for each site, highlighting active sites and their first neighbors using bolder points. While both *M*_S_ and *M*_SA_ reproduce well the observed *K − RSA, K − WCN*, and *K − B* trends, *M*_SA_ does a somewhat better job, reproducing observed trends better and, especially, showing a similar dispersion of rates around the smooth fits (e.g. sites near the active site are far from the trend both for observed and *M*_SA_ rates, but not for the *M*_S_ case). Regarding distance to the active site, while *M*_S_ fails to reproduce the correct *K − d*_active_ dependence, *M*_SA_ provides a good fit. Adding activity constraints is key to explain the dependence of site-specific rates on all the molecular traits that correlate with them.

In addition to smooth trends, Figure 3 shows the rates for each site. We see that *M*_SA_reproduces better than *M*_S_ not only the average trend, but also the *dispersion* of rates around the trend for *RSA, WCN*, and *B*: 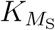 is much less variable around the smooth trend than 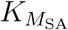 and *K*_obs_. The reason for this is that activity constraints, which are not accounted for by *M*_S_, contribute to this dispersion. For instance, active residues and their immediate neighbors deviate strongly from the smooth trends 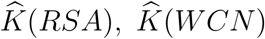, and 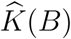 for both 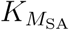 and *K*_obs_, but not for 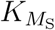. Thus, including activity constraints affects the rate-trait dependence not only for *d*_active_, where it was to be expected, but for all four of the molecular traits.

To test whether *M*_SA_ reproduces rate-predictor trends better than *M*_S_ for other proteins as well, I repeated the previous analysis on each of the 160 enzymes of the dataset. Figure 4 shows *RMSE*(*M*_SA_) vs *RMSE*(*M*_S_) for all proteins. Points below the diagonal represent cases for which *M*_SA_ reproduces trends better than *M*_S_. Counting cases for which Δ*RMSE* = *RMSE*(*M*_SA_) *- RMSE*(*M*_S_) *<* 0, we see that *M*_SA_ reproduces better than *M*_S_the dependence of site-specific rates with *d*_active_, *B*, and *WCN*, while it preforms similarly for the rate-*RSA* dependence (Table 1). Taking into account the magnitude of the improvements, as shown in Table 2, the average error difference (Δ*RMSE)* is negative for all four traits; *M*_SA_ fits rate-predictor trends better for all traits, with the largest improvement for *d*_active_ and the smallest for *RSA*.

**TABLE 1.**
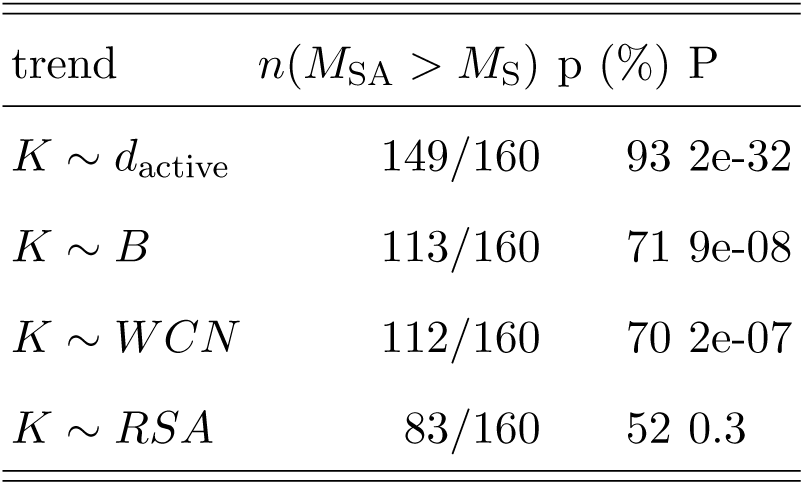
Binomial test of whether *M*_SA_ fits rate-trait trends better than *M*_S_. *n*(*M*_SA_ *> MS*) is the number of cases for which *M*_SA_ fits observed trends better than *M*_S_, *p* is the percentage of cases, *P* is the p-value.

| trend | $n(M_{\text{SA}} > M_{\text{S}})$ | p (%) | P |
| --- | --- | --- | --- |
| $K \sim d_{\text{active}}$ | 149/160 | 93 | 2e-32 |
| $K \sim B$ | 113/160 | 71 | 9e-08 |
| $K \sim WCN$ | 112/160 | 70 | 2e-07 |
| $K \sim RSA$ | 83/160 | 52 | 0.3 |

**TABLE 2.**
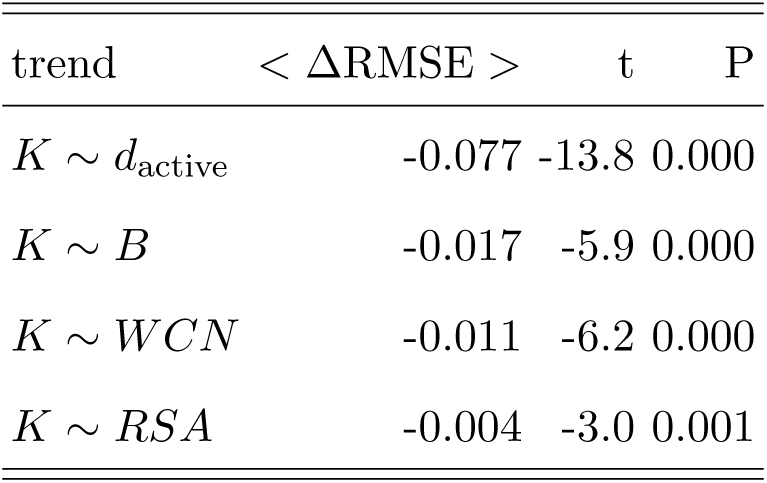
T test of whether *M*_SA_ fits rate-trait trends better than *M*_S_. ⟨Δ*RMSE*⟩ average over proteins of the difference between the *M*_SA_ and *M*_S_ errors of Figure. 4. Δ*RMSE* is negative (positive) when *M*_SA_ (*M*_S_) fits observed trends better. *t* is the T-test statistic and *P* the p-value. On average, *M*_SA_ fits all trends better than *M*_S_.

**FIG. 4.**
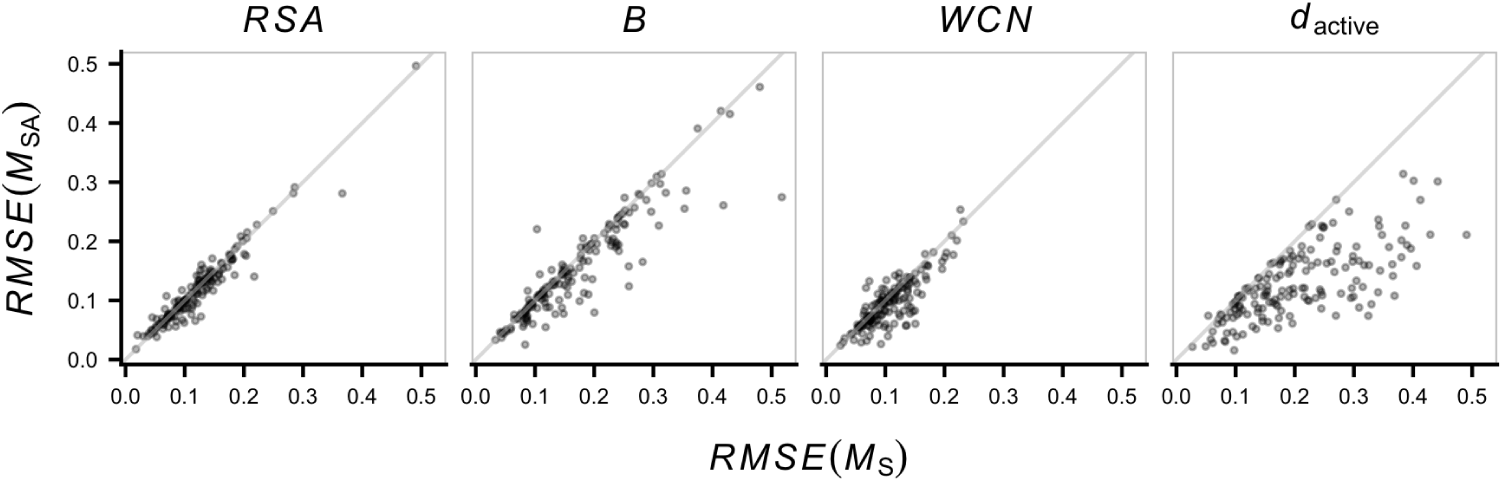
MSA provide reproduces rate-predictor trends better than MS. *RMSE*(*M*_SA_) (*RMSE*(*M*_S_)) is the Root Mean Square Error between the smooth rate-trend predicted by *M*_SA_ (*M*_S_) and the observed trend. Each point represents one protein. *RMSE* quantifies the fit between predicted and observed trends (lower is better; see Materials and Methods). For points below (above) the line *y* = *x*, which is shown for reference, *M*_SA_ provides a better (worse) account for the trend than *M*_S_. While, both models give similar *K ∼ RSA* trends, *M*_SA_ outperforms *M*_S_ in accounting for the *K ∼ WCN* and *K ∼ B* dependences. Notably, *M*_SA_ provides a much better account of the *K ∼ d*_active_ dependence for most proteins.

To summarize, *M*_S_ fails to reproduce the observed rate-*d*_active_ dependence (it overestimates rates of sites close to the active site and overestimates rates of distant sites), whereas *M*_SA_ accounts well for this dependence. Even though both models give good descriptions of the rate-trait dependence for *RSA, WCN*, and *B, M*_SA_ fits these trends better.

## DISCUSSION

I have posed two models of enzyme evolution, a model with selection for stability, *M*_S_, and a model with selection for stability and activity *M*_SA_. That selection acts on stability is well established and is the key assumption of the vast majority of biophysical models of protein evolution proposed so far [13]. However, natural selection acts not only on protein stability but on enzyme activity parameters such as *k*_cat_*/K*_*M*_ [1, 2]. Further, activation-energy factors are needed to explain, for instance, the effect of temperature on the growth rate, thus fitness, of viruses and bacteria [8, 20, 21]. Moreover, in some cases selection on activity is stronger than selection on stability [31]. Despite the extensive evidence of selection on protein activity, there is a general lack of biophysical models of evolution that consider activity constraints explicitly [13]. The *M*_SA_ model proposed here is a step towards filling this gap.

The key to *M*_S_, which was derived in detail previously [6, 14], is a fixation probability function that depends only on mutational stability changes, ΔΔ*G* (Eq. 1). In contrast, the fixation probability of *M*_SA_ depends not only on ΔΔ*G*, but also on mutational changes of the activation free energy of the enzyme-catalyzed reaction, ΔΔ*G*^*\**^. In [14], ΔΔ*G* was calculated using the all-atom empirical energy function of the program FoldX. Here, I calculated ΔΔ*G* (and ΔΔ*G*^*\**^) using a simpler coarse-grained Linearly-Forced Elastic Network Model (LFENM) Despite its simplicity, LFENM predicts ΔΔ*G* values with similar properties to those obtained using FoldX. (For instance, the average ΔΔ*G* increases with the number of contacts of a site. I will present these results in a separate publication.) More importantly, the models are validated by their success in fitting the empirical site-specific substitution rates and capturing the dependence of rates on several molecular traits.

Even if both *M*_S_ and *M*_SA_ predict site-specific evolutionary rates in good agreement with observations, I have found that *M*_SA_ fits observed empirical rates better. This means that evolutionary rate variation among sites in enzymes is determined not only but selection on thermodynamic stability, but also by selection on specific enzymatic activity.

The most noticeable difference between the two models is that *M*_SA_ reproduces correctly the dependence of site-specific rates on distance from the active site, *d*_active_, which *M*_S_ cannot explain. Thus, *M*_SA_ provides the mechanistic biophysical underpinning for the long-range influence of active sites on the evolution of other sites observed in phenomenological data analysis studies [10, 26].

Surprisingly, I found that activity constraints affect the dependence of rates on other molecular traits as well. *M*_S_ reproduces quite well the dependence of rates on *RSA, WCN*, and *B*, consistently with previous work that explained this dependence in terms of selection for stability [13, 15, 25, 27, 35]. However, *M*_SA_ fits the dependence of rates on these traits even better. Specifically, *M*_S_ tends to overestimate rates in regions of high *WCN* and low *B*. The reason is simple: active sites tend to be in these rigid (low *B*) highly-packed (large *WCN*) buried (low *RSA*) regions, where stability constraints are high, inducing extra activity constraints.

To summarize, I posed the model *M*_SA_, a new biophysical model of enzyme evolution that considers explicitly selection for stability and activity, and I compared *M*_SA_ with a stability-based model *M*_S_ on a diverse dataset of monomeric enzymes. I found that sitespecific substitution rates predicted by the model *M*_SA_ fit observed rates better than rates predicted by *M*_S_. Moreover, the dependence of observed rates on the molecular traits *RSA, B, WCN*, and *d*_active_ is better accounted for by *M*_SA_ than by *M*_S_. Thus, activity constraints have a significant effect on patterns of rate variation among sites and this is adequately modeled by the stability-activity model proposed here.

I believe the *M*_SA_ model will be helpful to explore several fundamental issues and to develop applications. For example, *M*_SA_ might be helpful to explain why proteins evolve to be moderately efficient [1, 2], just as stability-based models have explained why proteins are marginally stable [23, 57]. Another classical issue that may be explored using *M*_SA_ is the evolutionary implications of a trade-off between stability and activity [42]. On the applied side, *M*_SA_ could be used to improve active-site prediction [26]. Another application would be to add activity constraints, as modeled by *M*_SA_, to improve probabilistic evolution models used for phylogenetic inference purposes [49]. In general, I expect that developing biophysical models of protein evolution that consider selection for stability and activity, such as *M*_SA_, is a promising research endeavor that will advance our understanding of protein evolution and impact many areas of evolutionary biology.

## MATERIALS AND METHODS

### Two models of enzyme evolution

#### Origin-fixation models

When mutations are rare, evolution can be thought of as an origin-fixation process [38]. Most of the time the population consists of a single genotype, which is the current wild type. When a new mutant originates, it competes with the wild type and is destined to become either lost or fixed. For such an origin-fixation process, evolutionary dynamics is determined by the *fixation probability*, which is the probability that the mutant prevails, replacing the previous wild type. The two models of evolution described below are origin-fixation models.

#### The stability model (M_*S*_)

I consider a stability-based model, *M*_S_, that has been used to predict site-specific rates of evolution [14]. *M*_S_ is an origin-fixation model with fixation probability:

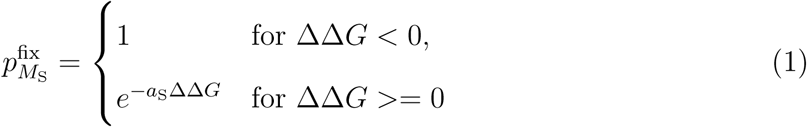

where ΔΔ*G* = Δ*G*_mut_ − Δ*G*_wt_ is the stability change due to the mutation. (Δ*G*_mut_ and Δ*G*_wt_ are the folding stabilities of mutant and wild type, respectively). The parameter *a*_S_ *≥* 0 represents the degree of selection pressure for protein stability.

#### The stability-activity model (M_SA_)

The purpose of this work is to go beyond stability constraints and study the effects of specific functional constraints on patterns of rate variation among sites. To this end, I pose a *stability-activity* model of enzyme evolution, *M*_SA_, with selection depending not only on stability but also on enzyme activity. *M*_SA_ is specified by the following fixation probability:

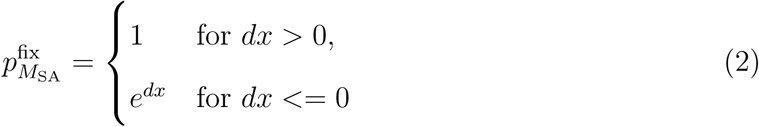

with

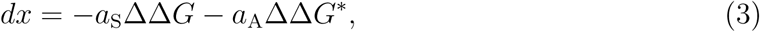

where ΔΔ*G* = Δ*G*_mut_ − Δ*G*_wt_ is the mutational change of stability and 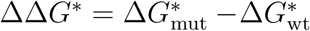 is the mutational change of the activation energy of the reaction catalyzed by the enzyme. The model has two positive parameters, *a*_S_ and *a*_A_, that represent the degree of selection pressure for stability and activity, respectively.

### Calculation of ΔΔ*G* and ΔΔ*G*^*\**^ using the Linearly Forced Elastic Network Model

To apply the models proposed above, we need to calculate mutational changes of stability, ΔΔ*G*, and of activation energy, ΔΔ*G*^*\**^. To this end, I use the Linearly Forced Elastic Network Model (LFENM) [11, 12, 25, 35].

#### Proteins are modeled as Elastic Networks

The LFENM models proteins as networks of nodes, representing amino acids, connected by harmonic springs, representing interactions. The energy function of a given wild-type protein is given by:

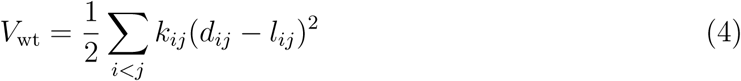

where *k*_*ij*_ and *l*_*ij*_ are, respectively, the force constant and equilibrium length of the spring connecting nodes *i* and *j*. Various Elastic Network Models have been developed over the years [4, 18, 34]. Here, I used a model proposed by Ming and Wall [43]: amino acids are represented by single nodes; nodes are connected if they are within *R*_0_ = 10.5*Å*; *l*_*ij*_ is the distance between the nodes in the PDB structure; the force constants are *k*_*ij*_ = 189 Kcal/Mol for sequence neighbors and *k*_*ij*_ = 4.5 Kcal/Mol otherwise. Since for evolutionary models it is better to represent residues using side chains rather than *C*_*α*_ [35], I placed nodes at the side-chain geometric centers (except for glycine, which has no side chain, for which I used *C*_*α*_ coordinates).

#### Mutations are modeled as perturbations

The LFENM models mutations as perturbations of the equilibrium lengths of the springs that connect the mutated site to its neighbors. Thus, introducing a mutation at site *r* results in a mutant with potential energy:

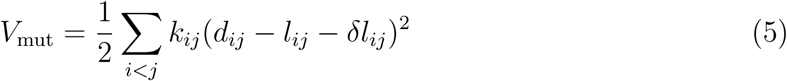

where *δl*_*ij*_ is a random perturbation added to *l*_*ij*_. Only contacts of site *r* are perturbed (i.e. *δlij ≠* 0 only for *i* = *r* or *j* = *r*, and *δl*_*ij*_ = 0 otherwise). I picked the non-zero *δl*_*ij*_ for the perturbed contacts independently from a normal distribution *N* (*µ* = 0, *σ = 0.3Å).*

#### Mutations distort the native conformation

*V*_mut_ is minimum at the mutant’s equilibrium conformation, which is given by:

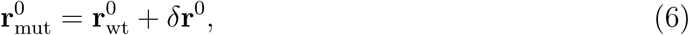

where 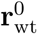 and 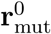 are the column vectors of node coordinates that represent the native conformations of the wild type and the mutant, respectively. The mutational conformational change can be calculated using:

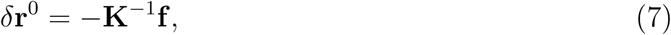

where **K** is the Hessian matrix of *V*_wt_ and *V*_mut_. (Since **K** is not affected by linear perturbations, **K**_mut_ = **K**_wt_ = **K**.) The vector **f** is a “force” that models the mutation and is calculated from the perturbations *δl*_*ij*_. For more details see [11, 12].

#### ΔΔ*G* and ΔΔ*G*^*\**^ are mechanical deformation energies

Using Eq. 4 and Eq. 5, assuming no variation of the unfolded state free energy, expanding potentials up to second order, and using basic statistical thermodynamics, it is possible to derive:

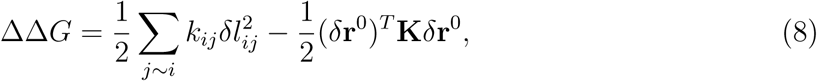

where ΔΔ*G* is the stability change due to the mutation, *δ***r**^0^ is the conformational change, calculated using Eq. 7, and *T* stands for transpose. The first term of Eq. 8 is always positive and represents the total stress introduced by the mutation [25, 35]; the second term is negative and represents the distortion of the mutant to relax part of the stress.

To calculate the change in activation free energy ΔΔ*G*^*\**^, I assume that the activation free energy is a sum of two terms a *distortion energy* plus a *vertical binding energy* [51, 56]. The distortion energy is the energy needed to change the conformation of enzyme and substrate from their free forms to their “poses” in the activated complex. The vertical binding energy is the free energy released when enzyme and substrate in the right poses bind together. I further assume that mutations at non-catalytic residues modify activity by distorting the active site’s structure, which modifies the distortion energy. Then, ΔΔ*G*^*\**^equals the energy necessary to deform the mutant’s catalytic residues so that they adopt their wild-type conformation, which can be derived from Eq. 5 ΔΔ*G*^*\**^ to be given by:

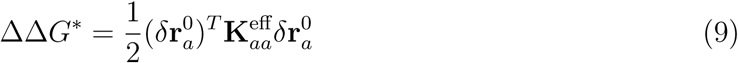

where 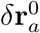 is the conformational change of the active residues due to the mutation and

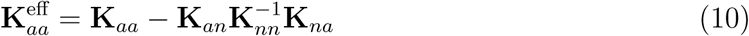

is a matrix that governs effective energies of deformation within the space defined by the coordinates of the active residues, taking into account their coupling with the rest of the enzyme. (To derive Eq. 10, the Hessian **K** was divided in four blocks:

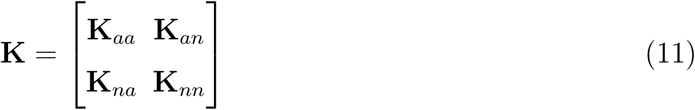

where the diagonal blocks **K**_*aa*_ and **K**_*nn*_ correspond to the active (*a*) and non-active (*n*) residues, respectively, and the off-diagonal blocks **K**_*an*_ and **K**_*na*_ couple active and non-active residues. See [41] for the derivation of an equation similar to Eq. 10.)

A note of caution is in order. Since the perturbation term of the LFENM is linear, the hessian **K** is not affected by the mutation, thus ΔΔ*G* (Eq. 8) and ΔΔ*G*^*\**^ (Eq. 9) do not include entropic changes that may have an effect on catalysis in the absence of conformational changes [39, 40].

### Testing models against data

#### Dataset

I tested the models on 160 of the 524 enzymes used in [26]. Specifically, I kept only monomeric enzymes. I further removed those enzymes that had missing amino acids or broken chains, which could result in wrong elastic network models. The dataset contains representatives of all main SCOP structural classes [44] and the six main EC functional classes [60]. No two enzymes of the dataset have more than 25% sequence identity. Catalytic residue information was obtained from the Catalytic Site Atlas [19]. PDB structures of these proteins were obtained from the RCSB protein database[5]. The list of proteins,their structural and functional classes, size and number of domains, and model parameters can be found in Supplementary Material file dataset.pdf.

#### Assessing whether models fit observed rates

First, I compared model rates with observed rates. Let *K*_obs_ denote the site-specific substitution rates estimated from multiple sequence alignments, which I call “observed” or “empirical” rates. I normalized these rates so that their average over sites is *< K*_obs_ *>*= 1: *K*_obs_ are rates relative to the protein mean. Briefly, calculating *K*_obs_ involves finding homologous sequences, aligning them, inferring a phylogeny, and using the sequence alignment and phylogeny as input for the program `rate4site` to estimate the substitution rate of each site [37]. Here I have used the site-specific rates calculated by Jack et al. [26], where the reader is referred to for further details.

Let *K*_*M*_ denote the site-specific rates predicted by model *M*, where *M* is either *M*_S_ or *M*_SA_. Let **a** be the list of parameters: **a** = *a*_S_ for *M*_S_ and **a** = (*a*_S_, *a*_A_) for *M*_SA_. The relative substitution rate of site *r* is given by:

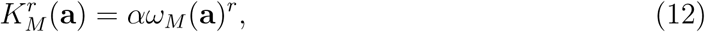

with

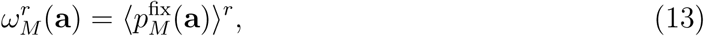

where 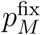 is the model’s fixation probability (Eq. 1 or Eq. 2), ⟨*…* ⟩ ^*r*^ stands for averaging over all possible point mutations at site *r*, and *α* is a constant that normalizes rates so that the average over sites is ⟨ *K*_*M*_ ⟩ = 1.

To calculate model rates for a given protein, I started by performing a full mutational scan: at each site, I introduced *N* = 19 simulated mutations and calculated, for each mutation, ΔΔ*G* (Eq. 8) and ΔΔ*G*^*\**^ (Eq 9). Then, for *M* = *M*_S_ and *M* = *M*_SA_, I determined the optimal parameter values minimizing the residual sum of squares 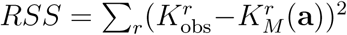 using the general purpose optimization function `optim` of R package **stats**.

To assess how well *M*_S_ and *M*_SA_ predictions fit the data, I compared model rates 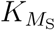 and 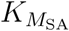 with observed rates *K*_obs_ for each protein. The most frequently used goodnessof-fit measure is the square Pearson correlation coefficient between model predictions and observations *R*^2^. Here, I used *R*^2^ to have an idea of the degree of improvement provided by *M*_SA_ over *M*_S_ (i.e. how much more of the variance of *K*_obs_ does *M*_SA_ account for). However, for non-linear models such as *M*_S_ and *M*_SA_, *R*^2^ is not adequate for model selection [55]. Thus, for model selection, I used the Akaike Information Criterion (*AIC*) and the associated Akaike weights *w*_*AIC*_ [55]. Specifically, for each model *M*, I calculated *AIC* = 2*k -* 2 ln *L*, where *L* is the maximum likelihood and *k* is the number of parameters. *AIC* is smaller for larger likelihoods and less parameters; the best model is that with the smallest *AIC*. Further, for each model *M*, I calculated the Akaike weight *w*_*AIC*_(*M*), which gives the weight of evidence for *M*, using 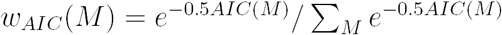.

#### Assessing whether models account for the dependence of rates on empirical predictors

In addition to assessing the *K*_*M*_ vs. *K*_obs_ goodness of fit, I studied whether the models are able to reproduce correctly the dependence of site-specific rates on molecular traits that characterize the local micro-environment of protein sites. Specifically, I considered four molecular traits that have been used as empirical predictors of site-specific rates: *RSA, WCN*, *B*, and *d*_active_, defined next.

- *RSA*^*r*^ is the relative solvent accessibility of site *r*. It measures the fraction of the site’s residue exposed to the solvent and is calculated as follows. First, the accessible surface area (ASA) is calculated using the software mkdssp [28, 29]. Then, ASAs are normalized by diving by the maximum solvent accessibility for each residue in a Gly-X-Gly tripeptide [58]. *RSA* = 0 is assigned to peptide linkages across chains, typically disulfide bridges. Here I used the *RSA* values of [26]. In principle, *RSA* should vary from *RSA* = 0 for completely buried sites to *RSA* = 1 for completely exposed sites. Sometimes, the calculation protocol results in *RSA >* 1, which is unphysical. In such cases, I set *RSA* = 1.
- *WCN*^*r*^ is the weighted contact number of site *r*, defined by:

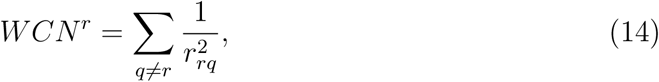

where, *r*_*rq*_ is the distance between the geometric centers of the side-chain atoms of residues *r* and *q*, respectively [35]. For glycine, which has no side chain, I used *C*_*α*_ coordinates.
- *B* is a measure that quantifies the local flexibility of a site. I calculated *B* using the B-factors available in the protein’s pdb file. The B-factor of an atom is proportional to its Mean Square Fluctuation (MSF) with respect to its coordinates in the native conformation. Here, I defined *B*, which I refer to as a residue B-factor, as the average of the B-factors of all heavy atoms of the residue’s side chain. For glycine, I used the B-factor of its *α* carbon instead.
- 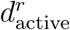 is the distance from residue *r* to the nearest catalytic residue. As for *WCN*, I considered distances between side-chain geometric centers (*C*_*α*_ for glycines). By definition, a residue is a catalytic residue if and only if *d*_active_ = 0.

To describe the dependence of site-specific rate *K* on site-specific molecular trait *X*, I obtained smooth fits to the *K* vs. *X* scatterplots, using local polynomial regression using the function loess of R package **stats**. Specifically, for each protein, I calculated smooth functions 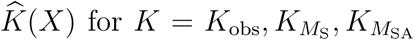, and *X* = *RSA, WCN, B, d*_active_. Then, I quantified how well *M* accounts for the *K*_obs_-*X* dependence using the Root Mean Square Error: 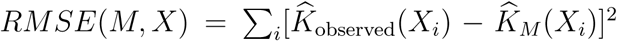, where *X*_*i*_ are evenly spaced points that cover the range of *X*. The model with lowest *RMSE*(*M, X*) is the one that best describes the *K*_obs_ *∼ X* dependence; Δ*RMSE*(*X*) *≡ RMSE*(*M*_SA_, *X*) *- RMSE*(*M*_S_, *X*) is negative (positive) if *M*_SA_ (*M*_S_) provides the best fit to the dependence of *K*_obs_ on trait *X*.

## ACKNOWLEDGMENTS

Most of this work was done at the Max Planck Institute for the Physics of Complex Systems (Dresden, Germany), where I was on sabbatical. This work was supported by Consejo Nacional de Investigaciones Científicas y T´ecnicas (grant number PIP 112 201501 00385 CO) and by Agencia Nacional de Promoción Científica y Tecnológica (grant number PICT-2016-4209).

## SUPPLEMENTARY MATERIAL

File dataset.pdf contains further information on the dataset.

